# Orchestrating Microbiome Analysis with Bioconductor

**DOI:** 10.1101/2025.10.29.685036

**Authors:** Tuomas Borman, Giulio Benedetti, Geraldson Muluh, Aura Raulo, Benjamin Valderrama, Artur Sannikov, Stefanie Peschel, Yihan Liu, Rasmus Hindström, OMA consortium, Katariina Pärnänen, Christian L. Müller, Aki S. Havulinna, Sudarshan Shetty, Marcel Ramos, Domenick J. Braccia, Héctor Corrada Bravo, Felix M. Ernst, Levi Waldron, Thomaz F. S. Bastiaanssen, Himel Mallick, Leo Lahti

## Abstract

The expansion of microbiome research has led to the accumulation of interlinked datasets encompassing versatile taxonomic and functional assays. The analysis of increasingly large and heterogeneous multi-modal microbiome data would benefit from unified approaches supporting the design of modular data science workflows through interoperable methods. The Bioconductor project has recently developed an optimized statistical programming framework for multi-assay data integration. Building on this foundation, we introduce a community-developed open source ecosystem for microbiome data science. In contrast to the previous alternatives, the methodology is specifically designed to support joint analysis of hierarchical, interlinked, and heterogeneous multi-table datasets that are increasingly common in modern microbiome research. This data science ecosystem encompasses open data, methods, tutorials, and an active online community. These resources support standardized and reproducible data wrangling, joint analysis, and reporting. We have detailed the functionality and usage in the online book https://microbiome.github.io/OMA, which offers guidance for prospective users and contributors.

## Introduction

Modern data science applications critically rely on open-source methods created by the research community ^1;2^. Open software ecosystems have become central innovation hubs, driving technological and cultural transformation ^3^. The Bioconductor project has become a significant distribution channel for open data science methods in the life sciences. Its vast developer community maintains over 2,300 quality-controlled R packages for various fields of life science informatics^4^. Additionally, Bioconductor has evolved into a global community that supports data science collaboration and training ^5^, playing an increasingly important role in advancing computational skills that are essential for modern microbiology education ^6^.

Microbiome research focuses on how microbial communities vary, function and interact with their environment and a potential host ^7^. The technologies used in this field generate vast amounts of high-dimensional and hierarchically structured data, commonly in the form of microbial DNA sequences detected in a set of samples. Following the rapid expansion of microbiome research and technological advances, metagenomic sequencing is increasingly complemented by other types of high-throughput measurements. This has led to the accumulation of multi-modal datasets—collections that combine different types of molecular data. These interlinked datasets encompasses diverse taxonomic and functional assays, ranging from microbial abundances and phylogenetic relationships to functional profiles.

Microbiome data scientists are thus increasingly facing the practical challenges of integrating heterogeneous data across multiple data modes into reproducible workflows. The unique statistical properties and complexity of microbiome data necessitate specialized approaches in their analysis ^7–9^, yet the diversity of proposed solutions and inconsistent outcomes ^10–12^ can make it difficult to identify the optimal methods and build interoperable workflows ^13;14^. Accordingly, a need for standardization has been highlighted ^7^. Open microbiome data science *frameworks* have emerged to provide sets of interoperable methodologies in specific computational environments ^15–18^, and a broader ecosystem of digital resources, vibrant user communities and collaboration culture that extend the capacities far beyond the underlying technical framework ^19^. However, frameworks focused solely on microbiome data science fall short on addressing the need to integrate data across diverse modalities, such as metagenomics, metabolomics, and host genomics.

The Bioconductor project embeds the methods of microbiome data science into a wider statistical programming framework alongside the concurrent development of interoperable methods and workflows for transcriptomics^20^, single-cell ^21^, metabolomics ^22^, proteomics ^22;23^, and other fields. Here, we introduce a mature data science ecosystem supporting the integration and analysis of multi-modal datasets in microbiome research. It provides harmonized data import and presentation, and fast, robust methods for transforming abundance tables into biological insights. The methods are extensively tested and distributed through R/Bioconductor packages (https://bioconductor.org/packages), supporting the community-driven development of the framework. To promote evidence-based best practices, we present the online book https://microbiome.github.io/OMA that introduces many dedicated topics in microbiome analysis.

## Data infrastructure

Data scientists routinely deal with interlinked assays, hierarchical data (*e.g.*, ontologies, trees), relational data (*e.g.*, networks), and supporting side information. This side information can include sample attributes like treatment group, collection site, or time point. The ability to integrate heterogeneous data elements is essential in biological research, where the individual components of a system can seldom be studied in isolation. However, input data formats vary across different methods, leading to technical complications and time lost on data wrangling. Bioconductor addresses this through standardized data structures supported by an ecosystem of interoperable methods, allowing the analyst to invest more time in core analytical tasks.

Bioconductor data structures support statistical analysis of complex data combinations ^24^ and are widely adopted by various research communities. They have been implemented through specialized object-oriented classes that integrate multiple data elements into a single structure known as a *data container* ^25^. The SummarizedExperiment (SE) class is the primary data container underlying the Bioconductor ecosystem ^24;26^.

Its position among the top 1% most-downloaded packages in Bioconductor reflects its widespread recognition and increasing adoption among developers (Figure 1). SE provides a standardized solution to store tabular data by linking numeric abundance matrices with side information on features or rows (*e.g.*, taxonomy), and samples or columns (*e.g.*, origin, collection time, host health).

**Figure 1:**
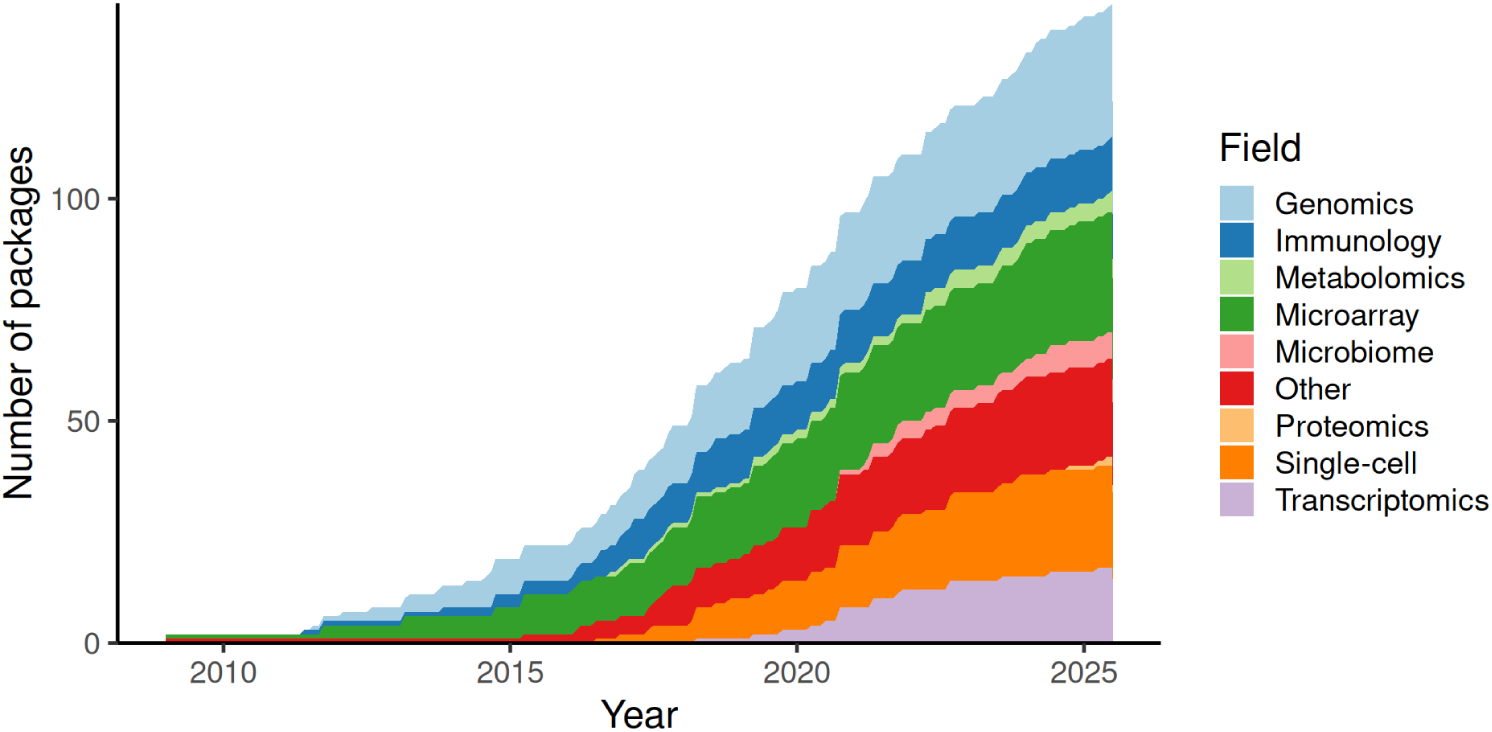
Adoption of the SummarizedExperiment (SE) data container among Bioconductor developers. The growth in the number of Bioconductor packages supporting SE over time is shown, categorized by their respective fields. Packages are counted by the date they were added to Bioconductor. Some packages added the SE support after their original release.

The interoperability of SE with other Bioconductor data structures facilitates the transfer of methods across different fields of omics research, yielding improved overall quality control and user support. This is a significant strength, which has facilitated the development of modular and scalable data science methods ^5^. Consequently, the SE-based framework has been extended to similar data integration tasks, notably in single-cell and spatial omics through the SingleCellExperiment (SCE) ^21^ and SpatialExperiment ^20^ classes.

### Hierarchical data structures

Microbiome data scientists frequently need to integrate information on feature phylogenies and sample hierarchies in order to accurately reflect fine-scale variability in microbiome data. While maintaining interoperability with the broader Bioconductor ecosystem, the TreeSummarizedExperiment (TreeSE) class ^27^ extends the SE and SCE data structures to hierarchical datasets by incorporating both feature and sample organizations as tree structures (Figure 3 a). Moreover, TreeSE supports the storage of other common aspects of microbiome data, for example, reference sequences, and incorporates results from common analytical procedures such as dimensionality reduction, statistical transformations, and agglomeration by taxonomic, functional or other information. The interoperability with other Bioconductor classes facilitates scalable analysis and the design of modular yet flexible workflows. In particular, the direct interoperability with the widely adopted SE data science ecosystem distinguishes our TreeSE-based approach from phyloseq, which is another popular Bioconductor data container developed initially with a focus on 16S amplicon data analysis ^18^. TreeSE can be seen as a more general framework than phyloseq, as it can accommodate all components typically included in a phyloseq object (taxa abundances, sample metadata, phylogenetic tree, and reference sequences). In addition, TreeSE can also accommodate multi-modal datasets encompassing parallel taxonomic levels, functional predictions, and resistome, metabolome or other profiles and alternative feature representations. It also offers substantial gains in memory efficiency and computing speed over phyloseq (Figure 2). While speedyseq is faster in certain operations, TreeSE tends to scale more efficiently in terms of execution time (t) and allocated memory (m) as sample size (n) increases. Despite these advantages, optimizing time and memory consumption presents an obvious area for further development in the TreeSE-based framework. Conversion functions and detailed cheat sheets between the two formats are available to support interoperability between the respective user communities.

**Figure 2:**
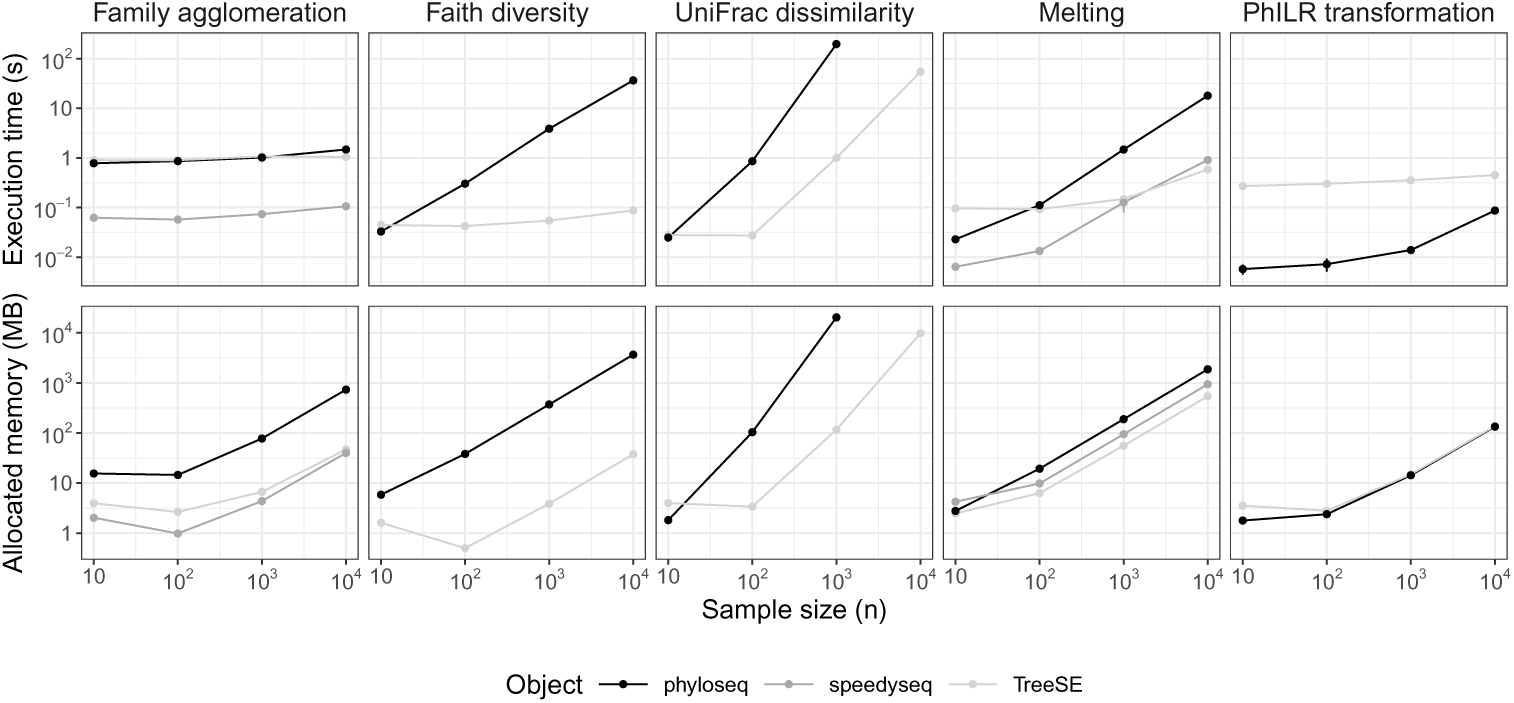
Execution time and memory consumption for microbiome data containers. The Bioconductor’s TreeSE, phyloseq and speedyseq are commonly used data containers in microbiome analysis. We benchmarked them for five common operations applied to random subsets of samples from a large study on wild baboons ^28^ (accessible through the microbiomeDataSets package in Bioconductor). Execution time (s) was measured as the mean value over 10 iterations, whereas allocated memory (MB) was estimated based on a single iteration. The benchmarking was conducted in R using the bench library with 8 CPU cores and 32 GB RAM. All operations, except for UniFrac dissimilarity with phyloseq, could also be performed on a regular laptop (*e.g.*, 4 CPU cores and 16 GB RAM). The source code for the benchmarking experiments is available through the OMA online book.

**Figure 3:**
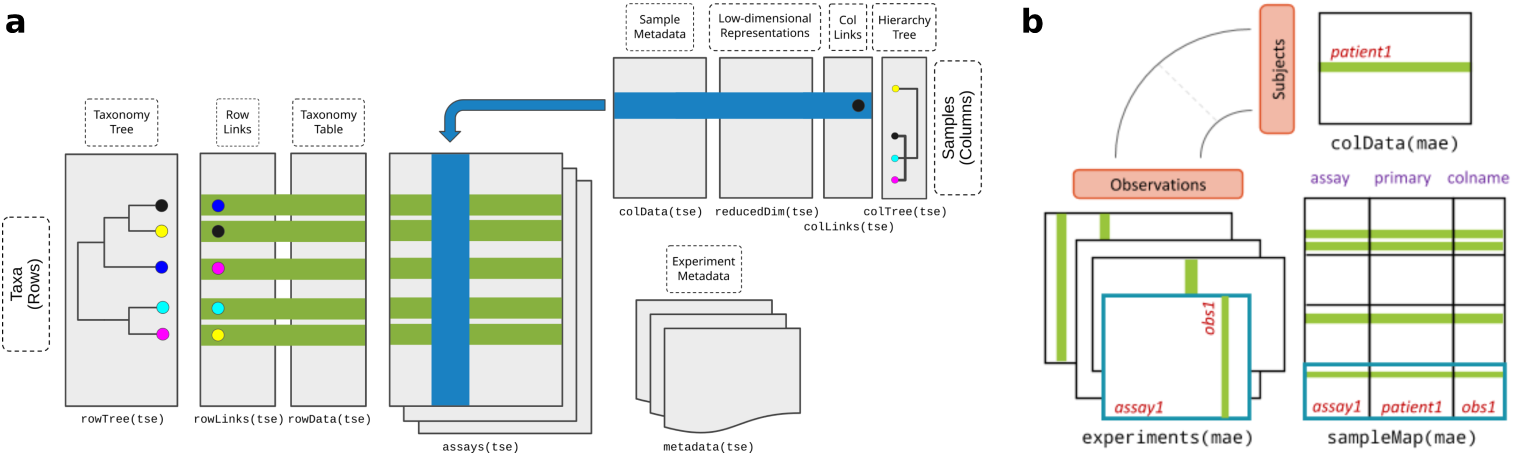
Bioconductor data containers for microbiome data. **(a) TreeSummarizedExperiment (TreeSE)** supports hierarchical, multi-table microbiome analysis. A data object can accommodate: i) “assays” that describe the abundances of features in each sample (*e.g.* taxonomic units, resistance genes, or predicted functions), ii) “colData” with tabular metadata on each sample, iii) “rowData” with tabular metadata on each feature (*e.g.* taxonomic mappings), and iv) tree information describing feature hierarchies, such as phylogenetic relations between the taxonomic units (rowTree) or host species (colTree). TreeSE can thus link multiple numeric abundance tables and their derived versions (*e.g.* statistical transformations, different taxonomic levels, dimensionality reduction) with tabular and hierarchical side information on the features (rows) and samples (columns). Additional slots for reference sequences and experiment metadata are available. (Figure adapted from ^27^) **(b) MultiAssayExperiment (MAE)** facilitates the integration of interlinked data sets that may represent different omics modalities linked by potentially non-trivial sample mappings; (figure from ^31^)

### Multi-assay data structures

Contemporary microbiome research frequently incorporates data across multiple measurement modalities for enhanced functional and mechanistic insights. This brings the need to find an appropriate representation for each data type and map the links between these. Whereas the TreeSE data container is broadly applicable to different modalities such as taxonomic, resistome, transcriptome, proteome, and metabolome profiling data, dedicated data containers have been developed for certain modalities by the Bioconductor community to address their specific characteristics; examples include single cell sequencing^21^ and host genomics ^29^. Subsequent linking of the data from distinct modalities often presents an additional challenge: multi-omics data is often sparse, so that samples can be only partially matched between experiments. In other cases, a single sample in one modality may link to two or more samples in another modality see ^30^. Integrating data across multiple modalities could thus benefit from flexible sample mappings and the ability to accommodate potentially different data containers corresponding to the different modalities. The MultiAssayExperiment (MAE) data container ^31^ provides a solution to these two tasks and facilitates the integration of different omics modalities (Figure 3 b). Integration of multi-table data without such a structured framework could lead to ambiguous mappings and inconsistent results. For instance, diagonal integration methods ^32;33^—where shared features across data types may not be explicitly defined—can benefit from such a systematic framework that enforces anchor alignment. This mirrors lessons learned from single-cell data integration, where shared anchors and hierarchical mapping have proven essential for interpretability. Flexible mapping of samples across several data objects supports efficient data integration, method sharing, and overall interoperability.

Table 1 summarizes the integrative data containers for microbiome analysis; Table 2 introduces the available data elements in each container.

**Table 1:**
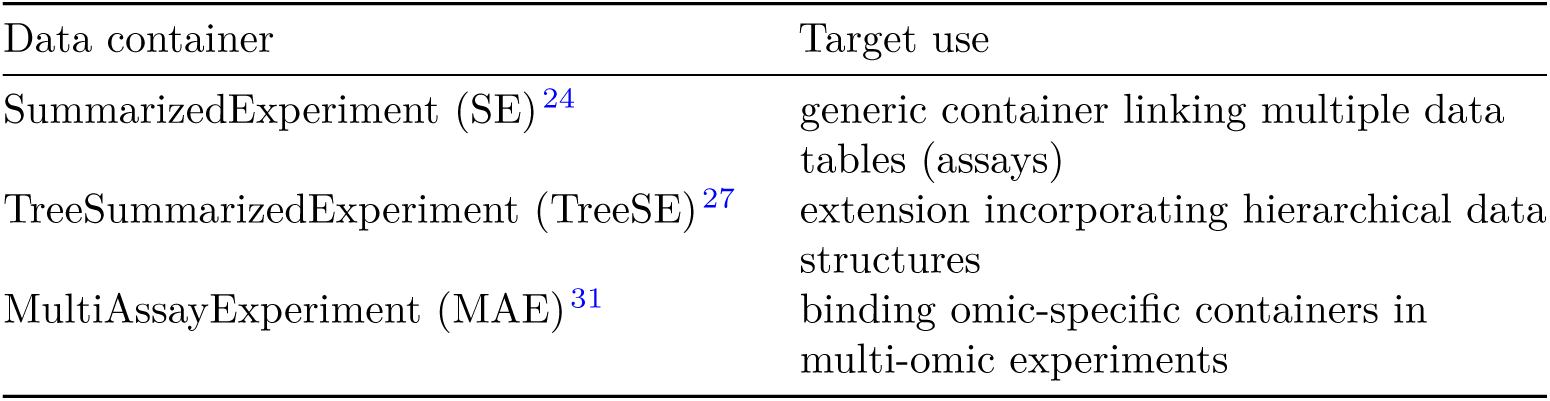
Integrative data containers for microbiome analysis.

| Data container | Target use |
| --- | --- |
| SummarizedExperiment (SE)<sup>24</sup> | generic container linking multiple data tables (assays) |
| TreeSummarizedExperiment (TreeSE)<sup>27</sup> | extension incorporating hierarchical data structures |
| MultiAssayExperiment (MAE)<sup>31</sup> | binding omic-specific containers in multi-omic experiments |

**Table 2:** Key data elements in the integrative data containers.

| Slot | Content | SE | TreeSE | MAE<sup>a</sup> |
| --- | --- | --- | --- | --- |
| **Tables** |  |  |  |  |
| assays | tables of features (rows) and samples (cols) | X | X | X |
| altExps<sup>b</sup> | experiments with variable number of features |  | X | X |
| reducedDims | results of dimensionality reduction |  | X | X |
| **Rows** |  |  |  |  |
| rowData | side information on features | X | X | X |
| rowPairs | linkage between pairs of associated features |  | X | X |
| rowTrees | hierarchical organization of features |  | X | X |
| referenceSeq | reference sequences of features |  | X | X |
| **Columns** |  |  |  |  |
| colData | side information on samples | X | X | X |
| colTrees | hierarchical organization of samples |  | X | X |
| sampleMap<sup>c</sup> | flexible sample mapping across experiments |  |  | X |
| **Other** |  |  |  |  |
| metadata | additional information on the experiment | X | X | X |

<sup>a</sup> While MAE does not contain conventional slots (with the exception of colData), its experiments can include any number of data containers that provide those slots. <sup>b</sup> altExp samples must match assay samples in a 1-to-1 mapping. <sup>c</sup> The sampleMap supports diverse mappings between samples from different experiments (1-to-1, 1-to-many, many-to-1 and many-to-many).

### Data resources

Several open microbiome data resources are supported, enabling direct data import into the aforementioned data containers for further downstream analysis within the Bioconductor methods ecosystem (see Table 3). Examples of the available data resources include curatedMetagenomicData^34^, HoloFood ^35^, microbiome-DataSets^36^, and the EBI/MGnify database ^37^. In addition, Bioconductor packages often include built-in demonstration datasets. Collectively, these resources encompass taxonomic and functional profiles from thousands of published microbiome studies and tens of thousands samples that can be accessed with the available tools for research and educational purposes.

**Table 3:** Supporting open microbiome data resources (accessed 29/09/2025).

| Data resource | Package | # studies<br>or datasets | # samples |
| --- | --- | --- | --- |
| Curated metagenomic data<sup>34</sup> | curatedMetagenomicData<sup>34</sup> | 93 | 22,588 |
| HoloFood<sup>35</sup> | HoloFoodR<sup>38</sup> | - | 9,990 |
| MGnify<sup>37</sup> | MGnifyR<sup>39</sup> | 5,129 | 629,821 |
| microbiomeDataSets<sup>36</sup> | microbiomeDataSets<sup>36</sup> | 6 | 19,100 |
| Microbiome<br>Benchmark Data<sup>12</sup> | MicrobiomeBenchmarkData<sup>12</sup> | 6 | 1,125 |
| Human Microbiome<br>Project 2<sup>40</sup> | HMP2data<sup>40</sup> | 3 | 11,493 |
| A compilation of open<br>microbiome data sets | https://github.com/microbiome/data | 8 | 18,777 |

### Data preparation

While microbiome data scientists continue to develop specialized solutions for data analysis ^8;9;41^, the implementations typically differ in their expected input and output formats. Common data standards support modular data science workflows, speeding up methods development, benchmarking, and replication. Building on these principles, we describe a set of interoperable software libraries that support microbiome data integration and analysis based on the two integrative data containers, (Tree)SE and MAE (Figure 4).

**Figure 4:**
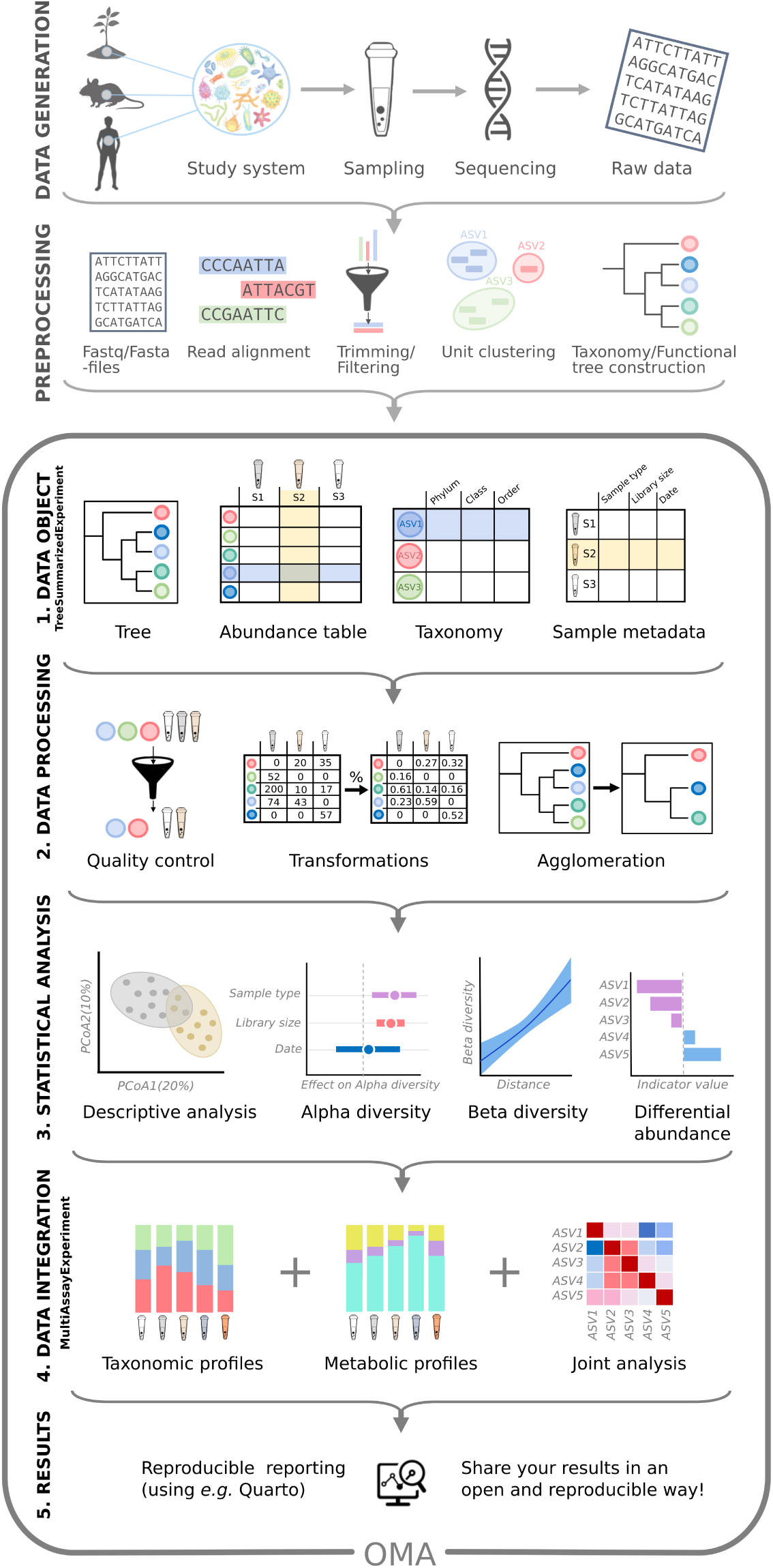
Multi-modal data integration workflow. Following data generation and preprocessing (top two panels), the integrative framework covers five steps of microbiome data integration and analysis: 1) Data is wrapped into a TreeSE data object (see Figure 3), containing information on the samples (tube icons) and their features (*e.g.* taxonomic, functional, or resistome profiles; colored circles). TreeSE objects can accommodate multiple assay tables as well as additional hierarchical structures. 2) The TreeSE data object can then be processed to get cleaner data. This can include filtering out low-quality samples and features, transforming assay data for example by normalizing it to enhance the comparability of sample-specific abundances, or agglomerating the feature data to specific taxonomic levels or functional categories. 3) The cleaned data can then be analyzed using statistical models to uncover relations among samples (*e.g.*, with PCoA), analyze drivers of variation in community diversity and composition or test for differential abundance (*e.g.* between sampling time or location). 4) The framework supports the integration of various types of omics modalities through MultiAssayExperiment and the specialized data structures for each omics provided by Bioconductor. Here, for example, the taxonomic and metabolomic profiles of samples can be summarized and compared with joint analysis among features. 5) Finally, Quarto notebooks can aid reproducible analysis and reporting.

The framework supports specifically the downstream analysis of high-throughput sequencing data from microbiome experiments after summarizing the original measurements into abundance tables (*e.g.*, taxonomic or functional profiles). Taxonomic abundance tables are one of the most commonly encountered data types. Common measurement platforms include amplicon-based marker gene studies (*e.g.*, 16S rRNA) and metagenomic sequencing, but alternative profiling techniques are available including phylogenetic microarrays ^42^ and high-density optical mapping ^43^. The interoperability of the data containers support modular workflow design for integrating different omics data into the broader Bioconductor ecosystem. This mitigates reproducibility challenges posed by rapidly evolving reference databases and profiling tools (*e.g.*, MetaPhlAn2 vs. MetaPhlAn4) by enabling structured coexistence and direct comparison within a unified, version-resilient framework ^44;45^.

#### Data import

The TreeSE data container can be populated with microbiome data from common tabular data types but non-standard file formats may require additional data wrangling. To streamline data import, we provide importers for a variety of standard microbiome file formats including BIOM^46^, MetaPhlAn/HUMAnN (MetaPhlAn v. 2–4, HUMAnN v. 3) ^47^, Mothur ^16^, QIIME2^17^ and the TAXonomic Profile Aggregation and standardization format (taxpasta) ^48^. These importers bridge the gap between the raw sequence processing and downstream statistical analysis in R, allowing researchers to focus on interpreting biological patterns rather than wrangling file formats. Moreover, converters between alternative data structures in R are available supporting seamless conversions between TreeSE and BIOM, DADA2 ^49^ and phyloseq (the mia::convert* functions), and TreeSE-based microbiome data objects can be exported into external services such as MicrobiomeDB ^50^ and in other languages (*e.g.*, Julia). These converters facilitate widespread access to microbiome analysis methods. Imported data sets from different omics modalities can then be interlinked within the MAE data container with its flexible sample mappings as described above. This integrated data object can then be analyzed with dedicated downstream methods.

#### Quality control and exploration

After importing the data into R, the recommended first step involves basic data exploration and quality control ^51^. In addition to methods for identifying contaminant sequences and calculating summary statistics, the framework includes a versatile set of custom visualization methods to assess data variability, outliers, homogeneity of variance and other aspects that may need to be considered in downstream data analysis. The mia and miaViz packages provide convenient access to many operations for the integrative data containers regarding common tasks in microbiome data exploration and visualization (Figure 5).

**Figure 5:**
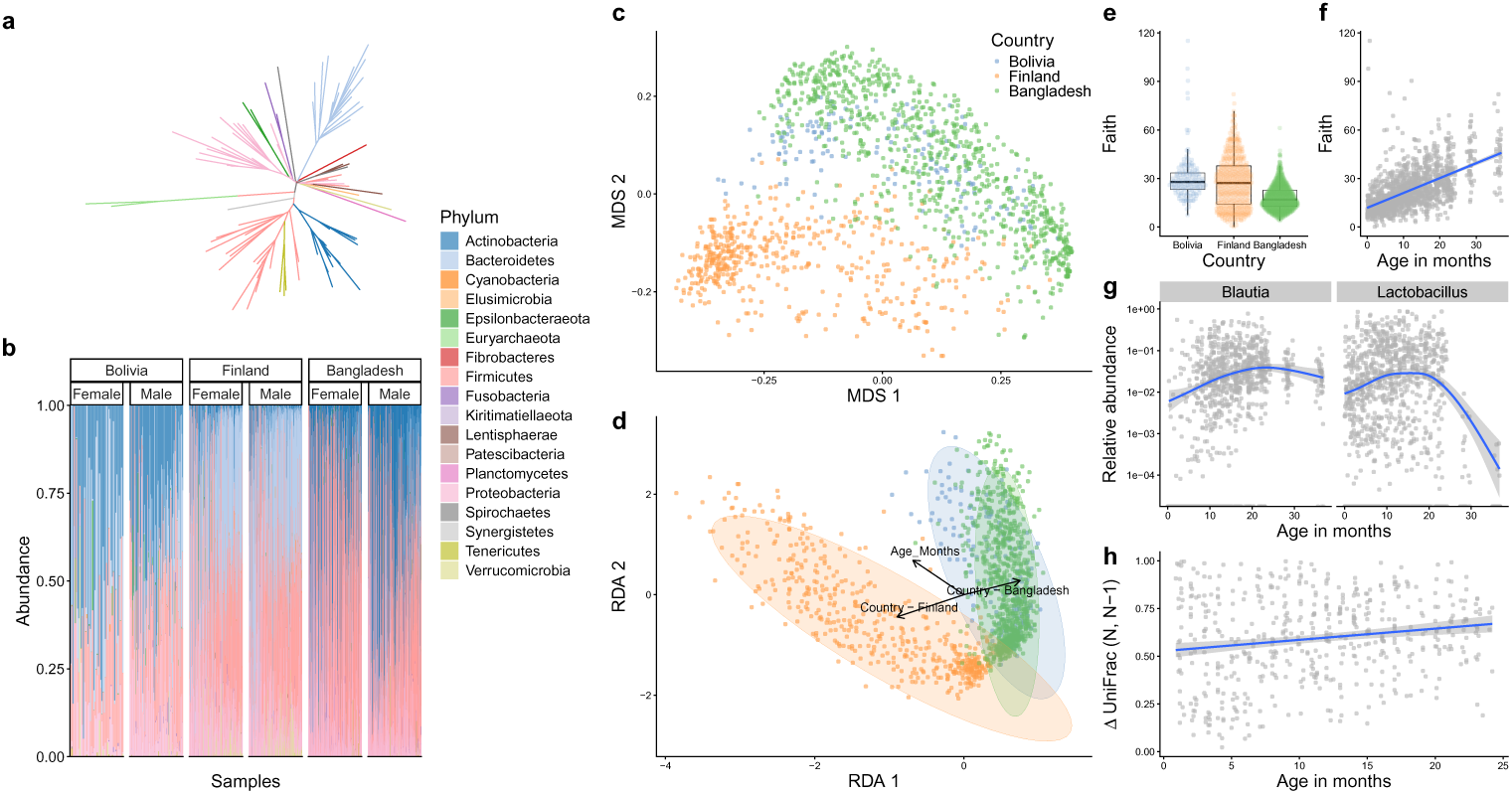
Common visualization methods applied to time series data on infants from three different countries (Finland, Bangladesh, and Bolivia) based on public datasets - SprockettTHData() ^97–99^ available via microbiomeDataSets^36^. **a**, Phylogenetic relationships. **b**, Phyla abundances of the samples ordered by age, stratified by country and gender. Actinobacteria is more abundant in Bolivia and Bangladesh compared to Finland. **c–d**, Multidimensional Scaling (MDS) using UniFrac distances and Distance-based Redundancy Analysis (dbRDA) based on Aitchison distance, respectively. Both ordination methods shows clear pattern separating microbial profiles in different countries. dbRDA shows an additional association between microbial profile and age. **e–f**, Faith’s phylogenetic diversity by country and age respectively. **g**, Differential Abundance Analysis (DAA) results. Two genera exhibit an association with age. **h**, UniFrac divergence between consecutive time points. Results indicate slow stabilization of microbial communities.

#### Filtering and agglomeration

Microbiome datasets can contain tens of thousands of unique features depending on the sequencing resolution, used reference databases, and the scope of analysis. Moreover, the integration of multiple omics modalities adds another level of challenges. To summarize data into biologically meaningful groups or to reduce multiple comparisons the data is often subsetted, filtered or agglomerated to higher-level taxonomic or functional categories or stratified by sample groups. The package ecosystem, and the mia package in particular, feature various tools for such tasks. For instance, the prevalence-based methods can be helpful to remove rare features that are likely sequencing artifacts or present otherwise limited value for the analysis. The mechanism of alternative experiments (altExp) allows agglomeration by taxonomic levels or other features such as pathogenicity or functional potential (*e.g.*, butyrate production ^52^, gut-brain modules ^53^, and other host-associated signatures ^54^).

The agglomerated data can be incorporated as a new alternative experiment within the original data object. The automated tracking of feature and sample links ensures that changes are propagated across all relevant elements of the data object, such as transformed abundance tables, phylogenetic trees, or interlinked omics modalities. This helps to avoid redundant processing and enables efficient management of interlinked multi-modal data sets.

#### Transformation

Statistical transformations commonly used in microbiome research include relative abundance (e.g. TSS) ^55^, tree-aware phylogenetic ILR (phILR)^56^, centered log-ratio (CLR), and robust centered log-ratio (rCLR). The latter three aim to mitigate compositionality bias ^57^. A related concept that controls for unequal sampling depth is data rarefaction. Rarefaction is the process of repeatedly subsampling data to equal read depth and averaging computed metrics ^58^. Subsampling data to equal read depth (*rarefying*) without iteration is not recommended ^59–61^. Our framework supports these and a variety of other transformations commonly applied to microbiome data.

#### Data enrichment

Incorporating additional data, for example, new versions of feature abundance data resulting from data transformations, is a frequent operation in applied data analysis, and it often takes place iteratively. The integrative data containers support gradual and organized accumulation of interlinked data elements including transformations, aggregated data, dimension reduction, and derived data on samples and features. We generally recommend initializing the data objects at the beginning of the workflow, where they can then be gradually enriched as the project advances. Data can be enriched by importing new external information as well as by deriving new information from the data itself into the same data container as described in the next section. By systematically extending the upstream data preparation step—instead of ad-hoc calculations scattered amidst the workflow—one can achieve a well-organized and transparent provenance of the integrated data object.

## Downstream data analysis

The framework is designed to improve the management of integrative analysis workflows. It facilitates joint operations on interlinked data elements, helping to shift the focus towards multi-modal analyses and methods development while also supporting standard data analyses in microbiome research.

Microbiome data analysis often relies on specialized statistical approaches since data sparsity, zero-inflation, compositionality, and heteroscedasticity pose challenges for standard methods ^8;55^. Moreover, microbiome data is hierarchically structured across taxonomic, ecological, temporal, and spatial levels ^7;62^. Alternative processing pipelines introduce different biases and generate varying data formats; for example, the same sample can yield differing taxonomic profiles depending on the computational method used ^63^. These and other unique characteristics have wide-reaching implications for microbiome analysis, necessitating dedicated analysis methods ^7^. The shared data containers support community-driven development and benchmarking of solutions. The following sections provide an overview of some of the currently available tools, encompassing standard methods for taxonomic and other abundance tables as well as integrative analyses between modalities.

### Community indicators

Various general indicators are available to quantify the microbial community state or other sample properties. Community diversity represents a common measure in microbial ecology ^64^ (Figure 5). Alpha diversity metrics generally quantify the observed or estimated number of distinct taxa (“richness”) and their distribution (“evenness”). Many diversity indices (*e.g.*, Shannon index), combine these two aspects. Phylogenetic diversity metrics additionally account the breadth of phylogenetic relatedness among community members ^65^, but are more computationally demanding than abundance-based metrics. However, a new implementation of the Stacked Faith’s Phylogenetic Diversity improves computational efficiency by 1–2 orders of magnitude ^66^. The framework supports analyses with rarefaction through the mia package ^58^. Other global indices of community state include measures of dominance, rarity, skewness, kurtosis, modality, library size (number of reads), or absolute abundance quantification. Sample information in the TreeSE object (colData) can be enriched with these derived quantities. By default, the mia::addAlpha method returns a set of complementary alpha diversity measures ^67^. Alpha diversity indices have been frequently applied to measure not only taxonomic diversity but also resistome diversity ^68^, and potentially other modalities.

**Community similarity** is commonly evaluated with beta diversity measures, which quantify multivariate dissimilarity between feature communities across all pairs of samples and can be used to compare community composition quantitatively. The framework supports common (dis-)similarity metrics in microbial ecology including Bray-Curtis, Jaccard, Aitchison ^57;64^, and the phylogeny-aware UniFrac ^69^. The community dissimilarities are commonly visualized with unsupervised ordination methods that project the data onto a lower-dimensional display ^70^. Among unsupervised techniques, those most frequently applied include Principal Component Analysis (PCA), Principal Coordinates Analysis (PCoA; also known as Multi-Dimensional Scaling, MDS), and Non-metric MDS (NMDS) (Figure 5). Statistical tests (*e.g.*, Permutational Multivariate Analysis of Variance) and supervised visualization methods, such as the distance-based redundancy analysis (dbRDA) can complement the unsupervised analyses, and help to uncover and quantify co-variation between community composition and sample-specific covariates. This can benefit from compositionally robust measures (*e.g.*, robust Aitchison distance ^57^), or phylogeny-aware dissimilarity measures specifically developed for microbiome research (*e.g.*, UniFrac ^69^). Sample dissimilarities provide the basis for many specialized microbiome techniques including microbial community typing (clustering) with Dirichlet Multinomial Mixtures (DMM)^71^ and community state types (CST)^72^, or factorization into community signatures ^73^. These and other multivariate methods are provided by mia, miaViz, and additional single-cell packages (scater, scuttle ^21^). Quantification of sample dissimilarities is a common task, and many omics modalities can benefit from access to the shared, standardized methods supporting the integrative data containers.

### Differential abundance and mediation analysis

Differential abundance analysis (DAA) and differential prevalence analysis (DPA) methods are used to identify indicator microbial features whose abundances vary across study groups or continuous gradients, for example, spatial or temporal distance between samples (Figure 5). Many dedicated DAA methods in microbiome research support the TreeSE framework, including methods such as ANCOM-BC2^74^, LimROTS^23^, MaAsLin3^75^, and radEmu ^76^. Many of these methods have also found applications with other omics modalities, such as transcriptomics (*e.g.* DESeq2), metabolomics ^75^, or proteomics ^23^ and the shared SEbased Bioconductor data containers have greatly facilitated this. Recent studies by us and others indicated that elementary methods, such as log transformed total sum scaled (TSS) counts coupled with a linear model, or presence/absence data with logistic regression often yield the most consistent results ^11;12^. The standard DAA typically focuses on individual features, ignoring interactions or other covariation among feature abundances. Thus, we support covariation analyses; indeed, agglomerating collinear features as a preprocessing step can reduce multiple testing, increase statistical power, and support interpretation, but is also prone to subjective parameterization and limited generalizability. The grouping can be based on clustering of the abundance profiles or on external information drawn from phylogenetic relations, taxonomic classifications or functional properties (*e.g.*, lump together all butyrate producers ^52^). Beyond these, first implementations of advanced methods—such as causal mediation analysis tailored to microbiome multi-omics data—have been implemented for SE^77^, highlighting its readiness to support the adoption of constantly developing methodology.

### Function- and sequence-level analyses

Taxonomic analysis is frequently complemented by functional analyses, either based on functional predictions derived from amplicon or metagenomic sequences, or from parallel metabolomic or other functional assays. Our framework not only supports the incorporation of functional abundance tables within the same data objects but specialized methods for functional data are available as well; for instance, regarding metabolome analyses, the notame package ^78^ has recently added SE support, and the R4MS project provides many interoperable tools ^22^. The framework has found recent applications in extended analyses of metagenomic features, for instance, antimicrobial resistome composition ^68^. We have also converted a large-scale compilation of resistome profiles in 8,972 metagenome samples ^79^ into the TreeSE format to support research and training activities in this area of microbiome analysis.

### Spatio-temporal, ecological, and epidemiological models

Microbiome time series ^80^ encompass a wide range of study designs, ranging from short to long follow-up times, sparse to dense data, univariate to multivariate and multi-omic monitoring^81^, and real, pseudo, or simulated time series. Moreover, an increasing number of studies incorporate spatial dimension in microbiome analysis ^62^. While spatio-temporal analyses require specialized methods, the miaTime R package offers a set of data wrangling methods to support longitudinal, multi-modal microbiome analysis. In addition, Bioconductor contains applicable methods from other research domains, namely single-cell sequencing ^21^ and spatial transcriptomics ^20^, which use closely related data containers. Spatial and temporal information can be incorporated as a covariate (*e.g.*, location or time) or by estimating temporally or spatially resolved metrics, such as divergence (dissimilarity between consecutive time points), autocorrelation (similarity of closely related samples), and feature modality (whether a feature exhibits multiple abundance states). Another type of temporal analysis includes the time-to-event, or survival models, which are encountered regularly in epidemiological and population cohort studies; for microbiome data, these analyses require specialized approaches, as proposed in previous studies ^82;83^. We also provide methods to simulate time series from standard models of microbial community dynamics including Hubbell’s neutral model, generalized Lotka-Volterra, and the consumer-resource model through the miaSim package ^84^. Our framework provides a robust basis for developing extended methods for longitudinal multi-omics, which is currently an active area of investigation ^85^.

### Multi-modal integration

A key benefit of the ecosystem lies in its ability to support the integration of microbial abundance data with other omics modalities, such as resistomes ^68^, metabolomes ^86^, host genomes ^87^, transcriptomes, or proteomes ^7;88^ (Figure 5). As we have demonstrated, the integrative data containers support complex combinations of omics data sets ^21;24;31^. Such integrative approaches in microbiome research can be broadly categorized into three groups: Machine Learning (ML) prediction ^89^, association-based approaches, and latent variable modeling ^7;8;85;89–91^. Examples of the currently supported methods for multi-modal analysis include cross-correlation, canonical correspondence analysis (CCA), anansi ^90^, and joint-RPCA^92^. Whereas the cross-correlation analysis can be used to quantify direct associations of samples or features between two data sets, anansi can supervise such analyses by incorporating known functional relations, and CCA and Joint-RPCA provide further tools to discover and visualize shared latent variation. We hope that systematic integrative frameworks, such as the one proposed here, will accelerate the development, benchmarking, and standardization of multi-modal extensions for microbiome analysis.

### Other methods and programming environments

A range of other methods have been employed in microbiome data science. Examples include network analysis^93^, ML-based prediction and classification ^94^, enrichment analyses via BugSigDB^54^ or Microbe Set Enrichment Analysis (MSEA) ^95^. The recent adoption of interoperable data containers and the methodological similarities in related fields can greatly expand the selection of available toolkits in emerging areas. In addition to the internal coherence within the Bioconductor ecosystem, this framework supports integration with external platforms through interfaces, namely reticulate for Python, Rcpp for C++, and compatibility with bioBakery pipelines ^47;96^ as well as Julia (MicrobiomeAnalysis.jl package). These connections empower users to combine the best tools available across environments and languages while maintaining structured data interoperability via standard containers such as TreeSE and MAE.

## Accessible analysis and community

Modern science is embodied by a critical community capable of assessing discoveries and replicating results ^100^. Open source developer communities present a timely manifestation of this principle, with technical, legal, sociological and epistemic consequences ^3^. FAIR principles for software support access to well-documented and user-friendly methods ^1^, and are facilitated by the extensive quality control and documentation standards enforced by Bioconductor ^24^. In addition to these technical demands, the development and quality control of academic software critically relies on active and critical communities that can provide continuous benchmarking and support the identification and dissemination of evidence-based best practices ^1;3;80^. We actively support community formation by online training resources and communication channels. Together with the interactive applications, these resources provide accessible entry points for all developers and users.

### The OMA book

*Orchestrating Microbiome Analysis with Bioconductor (OMA)* https://microbiome.github.io/OMA is an online book that provides a comprehensive introduction to the microbiome data science framework in Bioconductor. It offers practical guidance, reproducible workflows, and training material for integrative microbiome analyses. The book is versioned to match Bioconductor’s biannual software release cycles. The work incorporates rich feedback from the user community and microbiome training courses. It complements the broader trend, where specific Bioconductor data structures ^24;27;31^ have been adopted as a solution for multi-table data integration tasks in various domains such as single-cell sequencing (OSCA book) ^21^ and transciptomics (OSTA book). These resources demonstrate how standard data containers can support statistical programming on interlinked multi-table data sets. **Interactive applications.** Graphical user interfaces provide accessible entry points to the framework. The iSEEtree and miaDash packages provide interactive analysis and visualization tools for microbiome data ^101^. Their interface supports automatic generation of source code and replicable workflows as well as analysis templates for users without a strong programming expertise. Moreover, the tidy R paradigm, available through tidyomics, simplifies the syntax and can streamline the workflows ^102^. These tools allow users to analyze and visualize complex data combinations with ease.

### Community

Its widespread adoption has made R one of the most cited software resources of our time ^2^ and the introduction of the shared data structures facilitates the broad interoperability of the methods ecosystem. Bioconductor has a strong tradition in fostering the R user community through shared data structures and training activities ^5^. An active community of developers and users thrives online, supporting the software sustainability and promoting good practices in scientific computing ^103^. The community interacts via several online communication channels, including Bioconductor Zulip, GitHub platform, and an email list. The development of the framework is influenced by feedback from the user community through these channels, regular meetings, and training workshops.

## Outlook

In order to enjoy the benefits of automation while maintaining flexibility, data scientists routinely remix open-source software components into custom workflows. However, the lack of commonly adopted data representation standards often hinders the development, sharing, and application of modular, interoperable statistical workflows.

**The Bioconductor community** is widely recognized for developing open research software for life sciences. Its strengths include minimal documentation standards, portability across operating platforms, quality control (*e.g.* peer-review, unit testing, and continuous integration), usability (APIs), robustness (dependency testing and deprecation mechanisms), and a biannual release cycle with semantic versioning. Over the past two decades Bioconductor has grown into a repository of more than 2,300 actively maintained software packages, of which 62 are now classified under the *Microbiome* Task View. At the core of the Bioconductor’s data science strategy has been the idea of shared data representations ^4^ (Gentleman et al. 2004, @lawrence2013).

This has enhanced interoperability, mitigated redundant efforts and facilitated the translation of methods between research domains ^1;104^. In addition to these technical advances, the shared standards have played an important role in the formation of broad developer and user communities ^5^.

**Microbiome research** has shifted towards integrative analyses linking various omics’ towards enhanced mechanistic and causal insights ^64;105^. Technological advances in long-read sequencing, functional assays, spatial profiling, and single-cell analysis have created an increasing demand for integrative analysis techniques that can link microbiome data with other omics along varying spatial and temporal dimensions. In concert with parallel developments in other omics, our framework specifically addresses the need to integrate and jointly analyze interlinked abundance tables and supporting hierarchical and tabular side information across different omics. Whereas our framework supports the basic data wrangling and microbiomedata science operations, it also provides the structured input necessary for probabilistic inference and scalable computation. This is essential as the recent developments in microbiome and multi-omics workflows increasingly rely on advanced statistical and machine learning techniques such as Bayesian modeling and Gaussian processes to accommodate uncertainty and temporal or spatial structures ^106^. Building on these statistical foundations, the rapid emergence of foundation models and deep learning approaches in biomedicine calls for robust, scalable data representations ^107^ such as those we have adopted.

Applications include transfer learning across studies, interpretable embeddings of microbiome multi-omic profiles, computational prediction of microbial function and metabolites ^108;109^, and integration with AI agents for hypothesis generation and automated analysis, among others ^110;111^.

**Open data science infrastructures** have become critical components of reproducible research and a vast body of applications critically rely on open-source methods and workflows ^1;3;80^. Standardized data structures can enhance the transparency, reliability, and explainability of analysis workflows. We have documented the open data science ecosystem for multi-modal microbiome research built on widely adopted Bioconductor data containers, software libraries, data resources, tutorials, and a growing community of developers. Further extensions could include methods for temporal, spatial, and multi-omic analyses. Finally, supporting interoperability between R and other programming languages could facilitate the application of other machine learning frameworks. As the field evolves, the developer communities will continue to integrate new tools and approaches to improve integrative analyses of microbiome data.

## Consortia

OMA consortium: Bastiaanssen, Thomaz; Beber, Moritz E.; Bektanov, Aituar; Benedetti, Giulio; Borman, Tuomas; Braccia, Domenick J.; Calgaro, Matteo; Callan, Danielle; Chaudhary, Jiya; Corrada Bravo, Héctor; Courbayre Dussau, Basil; Das, Sneha; Eckermann, Henrik; Erawijantari, Pande; Ernst, Felix G.M.; Gaudron-Parry, Elliot; Gopalakrishnan, Shyam; Hakula, Maija; Hanhineva, Kati; Havulinna, Aki; Hindström, Rasmus; Jeba, Akewak; Lahti, Leo; Limborg, Morten; Lindgren, Himmi; Liu, Yihan; Mallick, Himel; Mongad, Dattatray; Müller, Christian L.; Muluh, Geraldson; Nynäs, Signe; Pasanen, Jesse; Paulo, Joao; Pelto, Juho; Peschel, Stefanie; Potbhare, Renuka; Pralas, Théotime; Ramos, Marcel; Raulo, Aura; Sannikov, Artur; Seraidarian, Ely; Serizay, Jacques; Shetty, Sudarshan; Shigdel, Rajesh; Suksi, Vilhelm; Tammi, Eetu; Valderamma, Benjamin; Waldron, Levi; Yang, Lu.

## Contributions

LL and TB initiated and coordinated the work. All authors contributed material, participated in manuscript preparation, and approved the final version.

### Acknowledgments

This project received funding from the European Union’s Horizon 2020 research and innovation programme under grant agreement No 952914. TB was additionally supported by the University of Turku Graduate School. We would like to thank the following individuals for their support for the project: Richa Ashma; Nitin Bayal; Chouaib Benchraka; Yang Cao; Axel Dagnaud; Teo Dallier; Noah de Gunst; Karoline Faust; Rob Finn; Samuel Hillman; Ruizhu Huang; Hervé Pagés; Lorna Richardson; Daniel Rios Garza; Mark Robinson; Matti Ruuskanen; Mahkameh Salehi. We would like to thank Christopher Quince and Kihyun Lee for providing the resistome profiling data, Laura Grieneisen and Johannes Björk for support with the baboon data compilation, Justin Silverman for philr support, and Ben Allen for MGnifyR support. We also extend our thanks to all contributors of the open-source resources used in this work and acknowledge CSC – IT Center for Science, Finland, for computational resources.

---

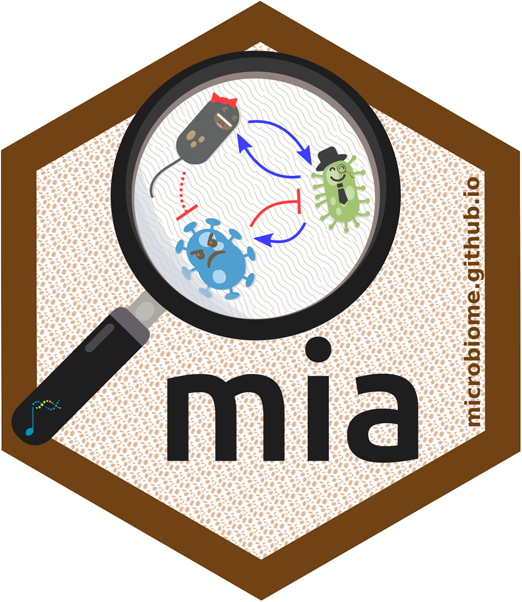

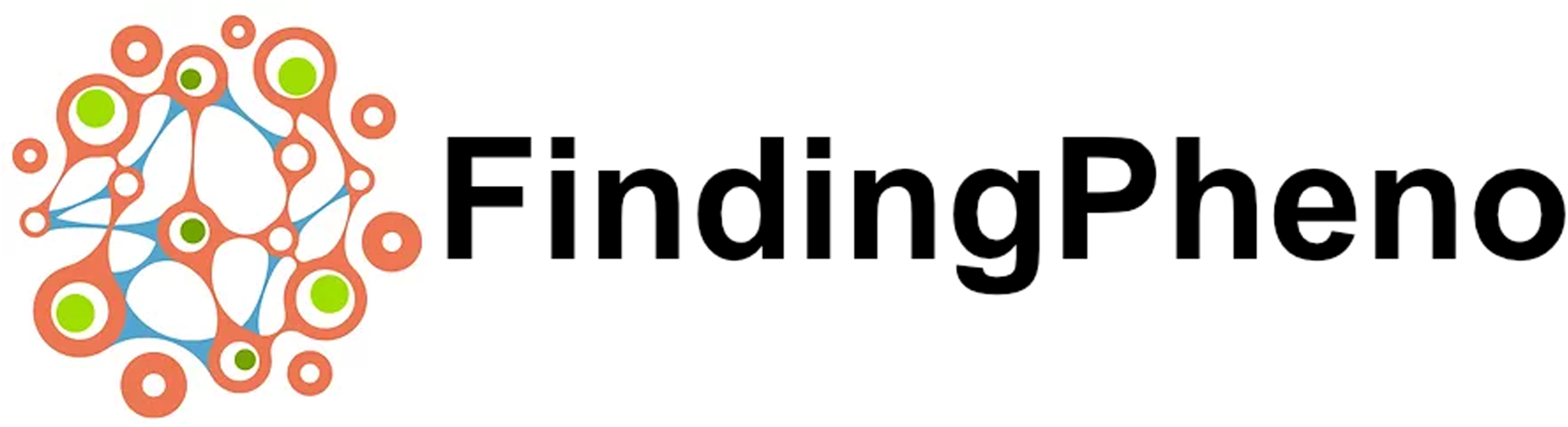

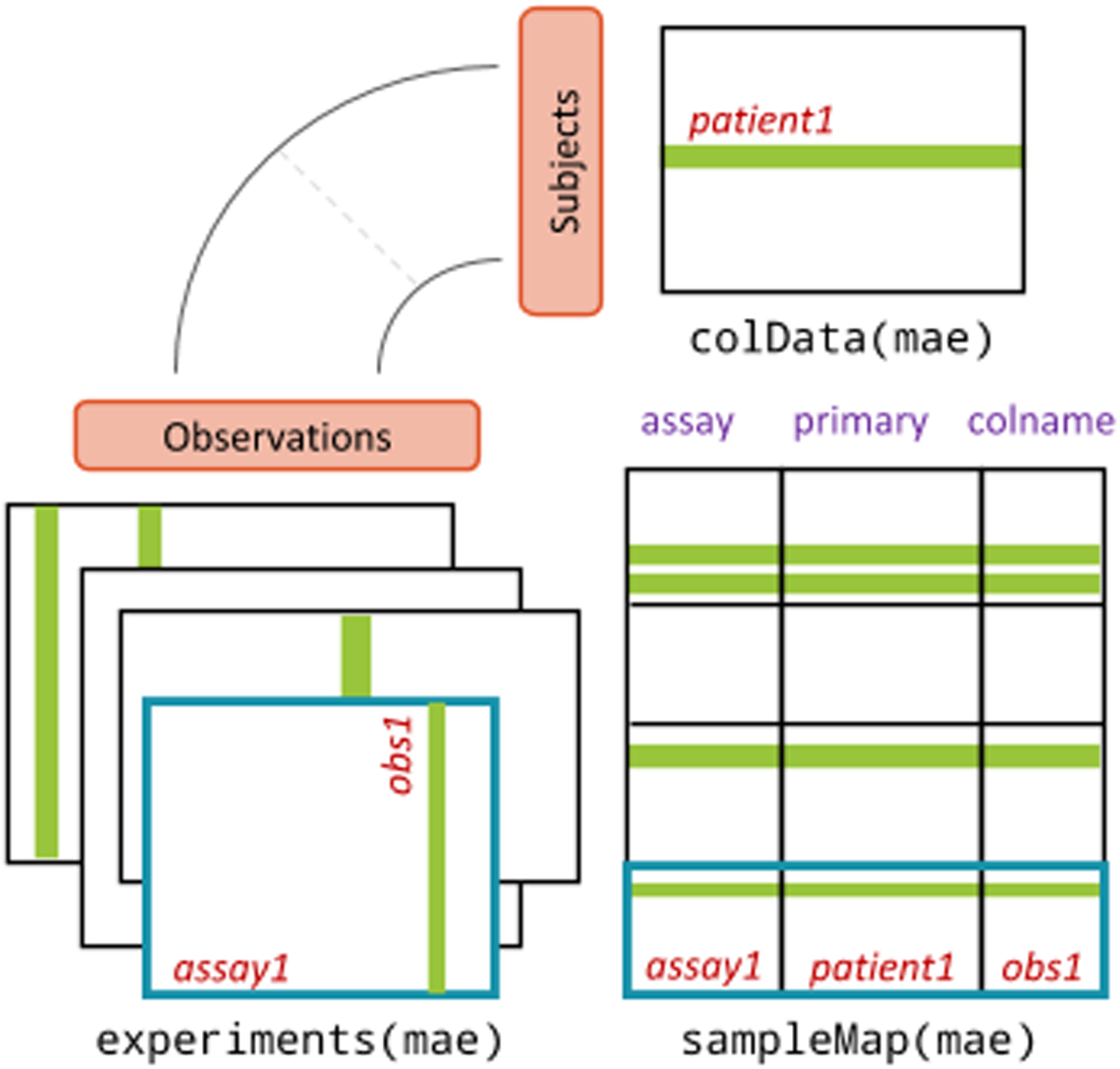

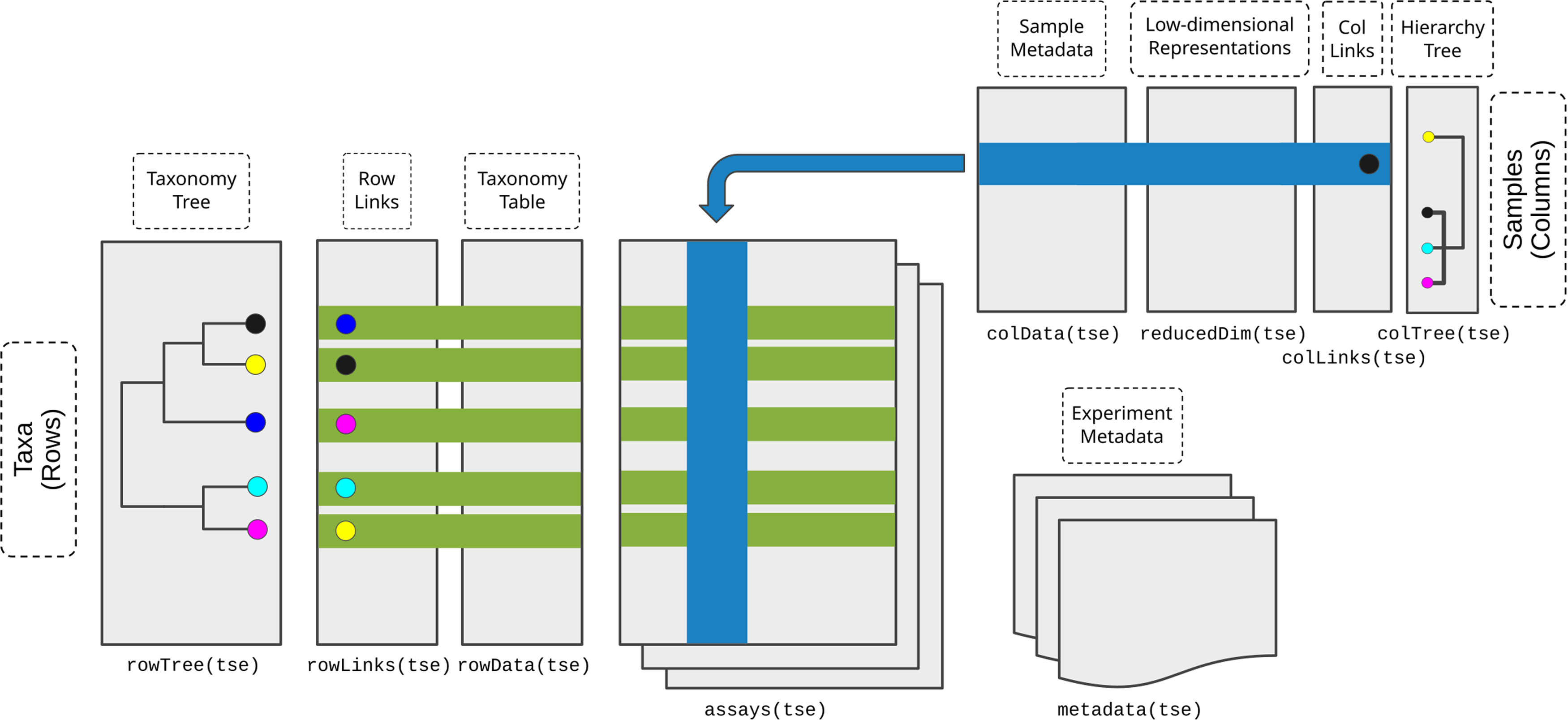

